# Activated BAX/BAK enable mitochondrial inner membrane permeabilisation and mtDNA release during cell death

**DOI:** 10.1101/272104

**Authors:** Joel S Riley, Giovanni Quarato, Jonathan Lopez, Jim O’Prey, Matthew Pearson, James Chapman, Hiromi Sesaki, Leo M Carlin, João F Passos, Ann P Wheeler, Andrew Oberst, Kevin M Ryan, Stephen WG Tait

## Abstract

During apoptosis, pro-apoptotic BAX and BAK are activated, causing mitochondrial outer membrane permeabilisation (MOMP), caspase activation and cell death. However, even in the absence of caspase activity, cells usually die following MOMP. Such caspase-independent cell death is accompanied by inflammation that requires mitochondrial DNA (mtDNA) activation of cGAS-STING signaling. Because the mitochondrial inner membrane is thought to remain intact during apoptosis, we sought to address how matrix mtDNA could activate the cytosolic cGAS-STING signaling pathway. Strikingly, using super-resolution imaging, we show that mtDNA is efficiently released from mitochondria following MOMP. In a temporal manner, we find that following MOMP, BAX/BAK-mediated mitochondrial outer membrane pores gradually widen over time. This allows extrusion of the mitochondrial inner membrane into the cytosol whereupon it permeablises allowing mtDNA release. Our data demonstrate that mitochondrial inner membrane permeabilisation can occur during cell death in a BAX/BAK-dependent manner. Importantly, by enabling the cytosolic release of mtDNA, inner membrane permeabilisation underpins the immunogenic effects of caspase-independent cell death.

## Introduction

To initiate cell death, apoptosis often requires mitochondrial outer membrane permeabilisation, or MOMP (Tait & Green, 2013). MOMP causes the release of mitochondrial intermembrane space proteins, including cytochrome *c*, that activate caspase proteases leading to apoptosis. Nevertheless, irrespective of caspase activity, cells typically do not survive following widespread MOMP, defining it as a point-of-no-return (Ekert, Read et al., 2004, Haraguchi, Torii et al., 2000, Tait, Ichim et al., 2014, Tait, Parsons et al., 2010). Because of this central role in dictating cell fate, MOMP is tightly controlled, primarily via pro-and anti-apoptotic members of the BCL-2 protein family (Lopez & Tait, 2015). Under homeostatic conditions, anti-apoptotic BCL-2 family members (e.g. BCL-2, BCL-xL, MCL-1) block the pro-apoptotic actions of BAX and BAK. Following an apoptotic trigger, BAX and BAK are activated, leading to their oligomerization in the mitochondrial outer membrane and MOMP (Cosentino & Garcia-Saez, 2017).

Mitochondrial apoptosis is considered a non-inflammatory form of cell death, allowing the host to quickly and efficiently clear away dead corpses without provoking an immune response (Arandjelovic & Ravichandran, 2015). Recently, others and ourselves have shown that caspase activity is essential for the non-inflammatory nature of mitochondrial apoptosis; if caspase activity is blocked following MOMP, cell death still occurs, but a type I interferon (IFN) response and NF-κB activation ensues (Giampazolias, Zunino et al., 2017#, Rongvaux, Jackson et al., 2014, White, McArthur et al., 2014). This leads to pro-inflammatory cytokine production and an immune response towards the dying cell that can initiate anti-tumour immunity (Giampazolias et al., 2017). Therefore, as proposed by others, a main function of apoptotic caspase activity may be to silence inflammation during cell death (Martin, Henry et al., 2012).

The ability of MOMP to activate inflammation and a type I IFN response requires recognition of mtDNA by the cytosolic cGAS-STING DNA sensing pathway (Giampazolias et al., 2017, Rongvaux et al., 2014, White et al., 2014). It is challenging to reconcile how matrix localised mtDNA can activate the cytosolic cGAS-STING DNA sensing pathway since the mitochondrial inner membrane is thought to remain intact during apoptosis (von Ahsen, Renken et al., 2000, Waterhouse, Goldstein et al., 2001). Given this paradox, we set out to define how mtDNA could activate cGAS-STING signaling. Using a variety of high-resolution imaging techniques, we find that following the onset of MOMP, BAX/BAK-mediated pores widen to allow the extrusion of newly unstructured inner membrane. Under conditions of caspase inhibition, the extruded inner membrane permeabilises facilitating mtDNA release into the cytoplasm, allowing it to activate cGAS-STING signaling and IFN synthesis.

Unexpectedly, our data demonstrates that the mitochondrial inner membrane can undergo permeabilisation during cell death. Importantly, by releasing mtDNA, mitochondrial inner membrane permeabilisation, or MIMP, enables cell death associated inflammation.

## Results

### mtDNA is released from mitochondria following MOMP

To visualise mtDNA dynamics during apoptosis, we used super-resolution Airyscan confocal microscopy. Importantly, this provides sufficient resolution to allow the visualization of the mitochondrial outer membrane and matrix-resident molecules (such as TFAM containing mtDNA nucleoids) as distinct entities under sub-diffraction-limiting conditions. As expected, in healthy U2OS cells, dual immuno-staining of the mitochondria outer membrane protein TOM20 and DNA revealed that mtDNA nucleoids are contained within mitochondria surrounded by a continuous outer membrane (**Figure 1A**). Next, to engage mitochondrial apoptosis, U2OS cells were treated with the BH3-mimetic ABT-737 (which inhibits BCL-xL and BCL-2 (Oltersdorf, Elmore et al., 2005)) together with actinomycin D (ActD), in the presence or absence of pan-caspase inhibitor qVD-OPh. To assess cell viability, U2OS cells were imaged using IncuCyte live-cell imaging and SYTOX Green uptake (**EV Figure 1A**). Consistent with engagement of mitochondrial apoptosis, ABT-737/ActD co-treatment rapidly and synchronously induced cell death in a caspase-dependent manner (**EV Figure 1A**). Under identical conditions, U2OS cells were immunostained with anti-TOM20 and DNA antibodies to visualise mitochondrial outer membrane integrity and mtDNA containing nucleoids respectively (**Figure 1B**). Specifically following ABT-737/ActD treatment, under caspase-inhibited conditions, permeabilisation of the mitochondrial outer membrane was observed, as evidenced by discontinuous, crescent-like TOM20 immunostaining (**Figure 1B**). Strikingly, mtDNA displayed cytosolic re-localisation in cells that had undergone MOMP under caspase-inhibited conditions (**Figure 1B**). A similar pattern of mtDNA cytosolic localization and mitochondrial outer membrane permeabilisation was also observed in E1A/Ras transformed MEF specifically following ABT-737/ActD/qVD-OPh treatment (**EV Figure 1B**). Given that both cytosolic and mitochondrial mtDNA structures were of similar size this suggests cytosolic re-localisation of mtDNA containing nucleoids. To determine whether mtDNA re-localisation was dependent on MOMP, we used CRISPR-Cas9 genome editing to delete BAX and BAK - two proteins essential for MOMP (**EV Figure 1C**)(Wei, Zong et al., 2001). In U2OS BAX/BAK^CRISPR^ cells, both MOMP and cell death were inhibited following ABT-737/ActD treatment, confirming effective functional deletion of BAX and BAK (**EV Figure 1D and 1E**). Under similar treatment, dual immuno-staining of TOM20 and DNA also revealed that BAX/BAK deletion prevented cytosolic release of mtDNA following ABT-737/ActD/qVD-OPh treatment (**Figure 1C**). We next investigated the dynamics of mtDNA re-localisation by live-cell imaging. The mitochondrial outer membrane was visualised using a SNAP-tag Omp25 fusion protein and the far-red fluorophore Janelia Fluor 646 (JF_646_) (referred to as JF_646_-MOM) (Grimm, English et al., 2015, Katajisto, Dohla et al., 2015). Because live-cell imaging of mtDNA using PicoGreen was not possible due to photobleaching, we monitored the localisation of mClover-fused TFAM (a mtDNA-binding protein present within mitochondrial nucleoids). As expected, TFAM-mClover localised to the mitochondrial matrix, co-localising with mtDNA (**EV Figure 1F**). U2OS cells expressing JF_646_-MOM and TFAM-mClover were treated ABT-737/ActD +/- qVD-OPh and imaged by live-cell imaging (**EV Figure 2**, **Videos 1, 2 and 3**). Under caspase-proficient conditions, following MOMP, the cell rapidly shrunk and detached, consistent with engagement of apoptosis (**Video 1**). Under caspase-inhibited conditions, after ABT-737/ActD treatment, TFAM-mClover was released into the cytosol, consistent with our earlier analysis of mtDNA (**Video 2**). Finally, we investigated the kinetics of TFAM-mClover release and MOM pore widening relative to MOMP. U2OS cells stably expressing TFAM-mClover, Omi-mCherry and JF_646_-MOM were treated with ABT-737 in combination with the MCL1 inhibitor S63845 in the presence of qVD-OPh then subject to live-cell imaging (**Figure 1D**, **Video 4**). Utilising this, in a sequential manner, mitochondrial release of Omi-mCherry (denoting MOMP) occurred prior to the visual appearance and widening of MOM pores (**Figure 1E**), finally followed by TFAM-mClover release. Collectively, these data show that matrix localised mtDNA and TFAM are released from mitochondria following BAX/BAK-dependent MOMP under caspase-inhibited conditions.

**Figure 1.**
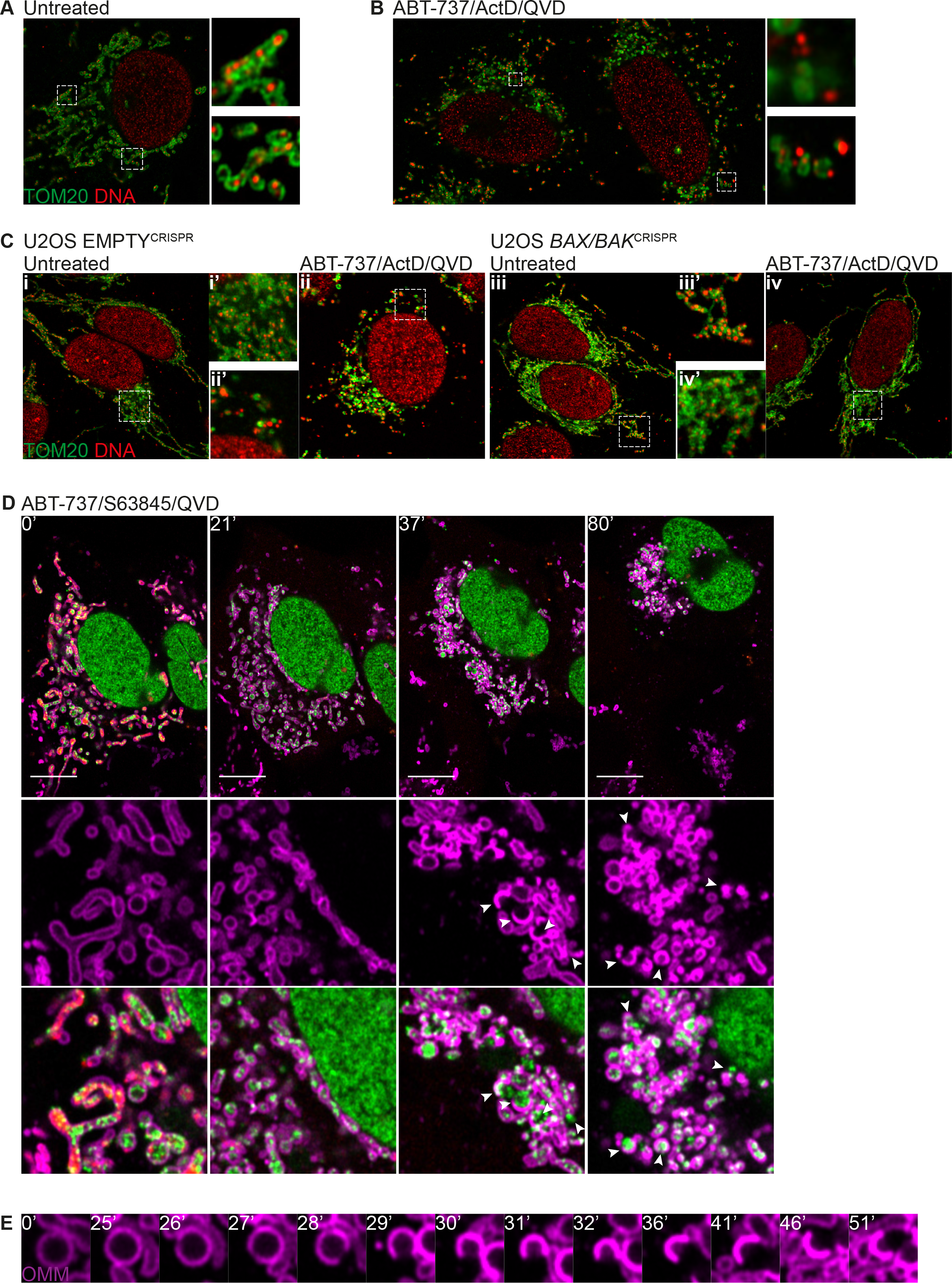
(**A**) Fixed super-resolution Airyscan images of U2OS cells immunostained with anti-TOM20 and anti-DNA antibodies. (**B**) Airyscan images of U2OS cells treated with 10 μM ABT-737, 1μM ActD and 20μM qVD-OPh for 3h, immunostained with anti-TOM20 and anti-DNA antibodies. (**C**) Airyscan images of control U2OS cells (EMPTY^CRISPR^) and U2OS cells with BAX and BAK deleted by CRISPR-Cas9 (BAX/BAK^CRISPR^) treated with 10μM ABT-737, 1μM ActD and 20μM qVD-OPh for 3h and immunostained with anti-TOM20 and anti-DNA antibodies. (**D**) Live-cell imaging of U2OS cells stably expressing JF_646_-MOM and Omi-mCherry, and transiently expressing TFAM-mClover. Cells were treated with 10μM ABT-737, 2μM S63845 and 20μM qVD-OPh at t=0. Scale bar: 10μm. (**E**) Zoom of a single mitochondrion from (**D**) and **Video 2** showing only the JF_646_-MOM. (**A, B, D, E**) are representative images of n≥2 independent experiments. (**C**) shows representative data from n=2 independent experiments.

### Mitochondrial inner membrane permeabilisation allows mtDNA release into the cytosol

Thus far, our data demonstrate that mtDNA/TFAM complexes relocalise beyond the mitochondrial outer membrane, however the integrity of the inner membrane (IMM) or localisation of mtDNA relative to the IMM was not determined. To investigate this, U2OS cells were treated with ABT-737/ActD in the presence of qVD-OPh, immunostained for apoptosis-inducing factor (AIF) and DNA (to visualise the mitochondrial IMM and mtDNA respectively) and analysed by Airyscan imaging (**Figure 2A**, **Videos 5 and 6**). In healthy cells the majority of mtDNA signal was within AIF immunostained structures, consistent with AIF being inner membrane localised and mtDNA residing within the matrix (**Figure 2A**, **Video 5**). Importantly, following MOMP (ABT-737/ActD/qVD-OPh treatment), mtDNA was found to localise in the cytoplasm, outside the inner membrane (AIF immunostained structures)(**Figure 2A**, **Video 6**). This suggests that mtDNA can pass beyond the inner membrane during caspase-independent cell death (CICD). To investigate this further, we simultaneously imaged the MOM, IMM and mtDNA during cell death. U2OS cells stably expressing JF_646_-MOM were treated with ABT-737/ActD/qVD-OPh and then immunostained with anti-AIF and anti-DNA antibodies to detect the MOM, IMM and mtDNA respectively. Cells were analysed by 3D-structured illumination microscopy (3D-SIM) and Airyscan imaging (**Figure 2B, EV Figures 3A and B**). 3D-SIM analysis demonstrated re-localisation of both AIF and mtDNA beyond the mitochondrial outer membrane specifically following MOMP (**EV Figure 3A**). Consistent with this, Airyscan analysis also demonstrated AIF and mtDNA relocalisation beyond the mitochondrial outer membrane specifically following MOMP (**Figure 2B**). 3D-Imaris based analysis revealed mtDNA and inner membrane protrusion through the mitochondrial outer membrane following MOMP (**Figure 2B and EV Figure 3A)**, in line with earlier live-cell analysis of TFAM-mClover, showing its extrusion (**Figure 1D**, **Videos 2 and 3**). In agreement with our earlier data, mtDNA also relocalised also outside the mitochondrial outer membrane following MOMP, either associated or not with the inner membrane (**EV Figures 3A and B**). Together, these data suggest that following MOMP, the mitochondrial inner membrane is extruded through the permeabilised outer membrane. Mitochondrial inner membrane permeabilisation, or MIMP, can then occur, allowing mtDNA egress into the cytosol.

**Figure 2.**
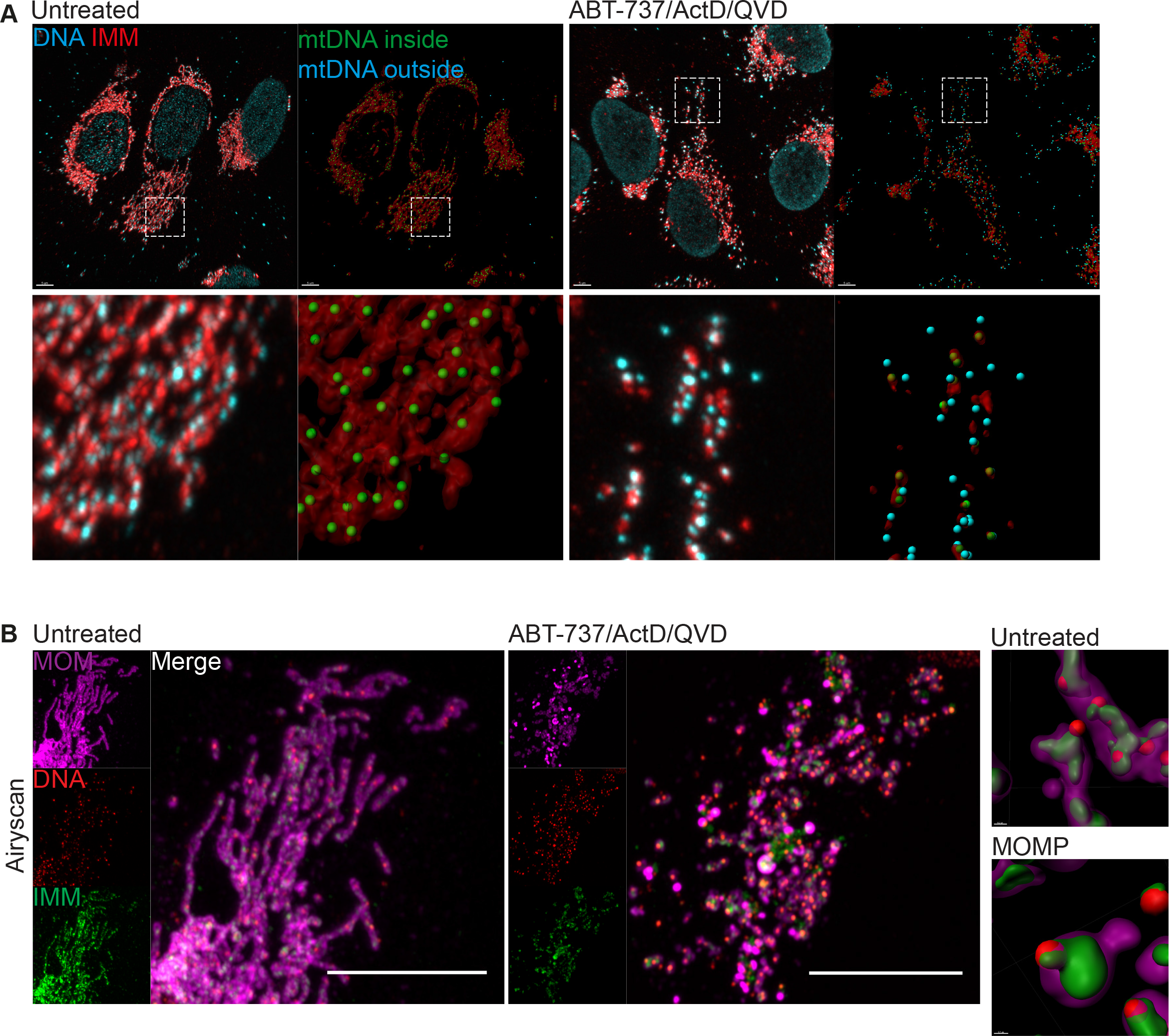
(**A**) 3D-view of Airyscan z-stack data of U2OS cells treated with 10μM ABT-737, 1μM ActD and 20μM qVD-OPh for 3h. Images were 3D rendered into surfaces for IMM (immunostained by AIF) and spots for DNA. Imaris analysis was performed to identify spots that lay within (green spots) and without (red spots) the surface. Scale bar: 5μm (**B**) Maximum intensity projections of z-stack Airyscan images of fixed U2OS cells stained prior to treatment with JF_646_ to label the MOM and immunostained after fixation for IMM (AIF) and DNA. Cells treated with 10μM ABT-737, 1μM ActD and 20μM qVD-OPh for 3h. Imaris 3D reconstructions of z-stack data shows the localisation of DNA and AIF in control and treated cells. Scale bar: 10μm. (**A-B**) are representative images of n≥3 independent experiments.

### Mitochondrial release of mtDNA occurs independently of mitochondrial dynamics and permeability transition

Various studies have shown that upon MOMP the GTPase DRP-1 promotes mitochondrial network fragmentation and inner membrane remodelling (Desagher & Martinou, 2000, Estaquier & Arnoult, 2007, Frank, Gaume et al., 2001). Potentially, inner membrane remodelling and/or mitochondrial fission at the point of MOMP might facilitate MIMP and mtDNA release. To investigate a role for DRP-1, we used conditional Drp1^fl/fl^ MEFs, using adenovirus expressing Cre recombinase to effectively delete DRP1 protein (**EV Figure 4A**) (Wakabayashi, Zhang et al., 2009). Wild-type or DRP-1 deleted MEF were then treated with ABT-737/ActD to engage mitochondrial apoptosis. Cells were imaged using IncuCyte live-cell imaging and SYTOX Green uptake. ABT-737/ActD treatment effectively induced apoptosis to a similar extent irrespective of DRP-1 expression (**EV Figure 4B**). In the presence of caspase inhibitor, samples were immunostained for TOM20 and mtDNA, and identified as having undergone MOMP by loss of cytochrome *c* staining. 3D-images generated were generated from Z-stacks of high-resolution images and quantified for mtDNA release (**Figures 3A** **and 3B**). In control samples, DRP-1 deleted cells displayed hyperfused mitochondria, as expected (**Figure 3A**). Importantly, the extent of mtDNA release in cells that had undergone MOMP failed to reveal a difference between wild-type and DRP-1 deficient cells (**Figures 3A** **and 3B**). To investigate a role for mitochondrial fission using an alternative method, we investigated if mitochondria that have already undergone fission could release mtDNA upon MOMP. To this end, U2OS MCL-1^CRISPR^ cells were pre-incubated with carbonyl cyanide *m*-chlorophenyl hydrazine (CCCP) to induce extensive mitochondrial fission, and then treated to undergo MOMP by addition of ABT-737 in the presence of qVD-OPh. Cells were immunostained for TOM20 and mtDNA then analysed by Airyscan microscopy. As expected, CCCP treatment induced extensive mitochondrial fragmentation (**Figure 3C**).

**Figure 3.**
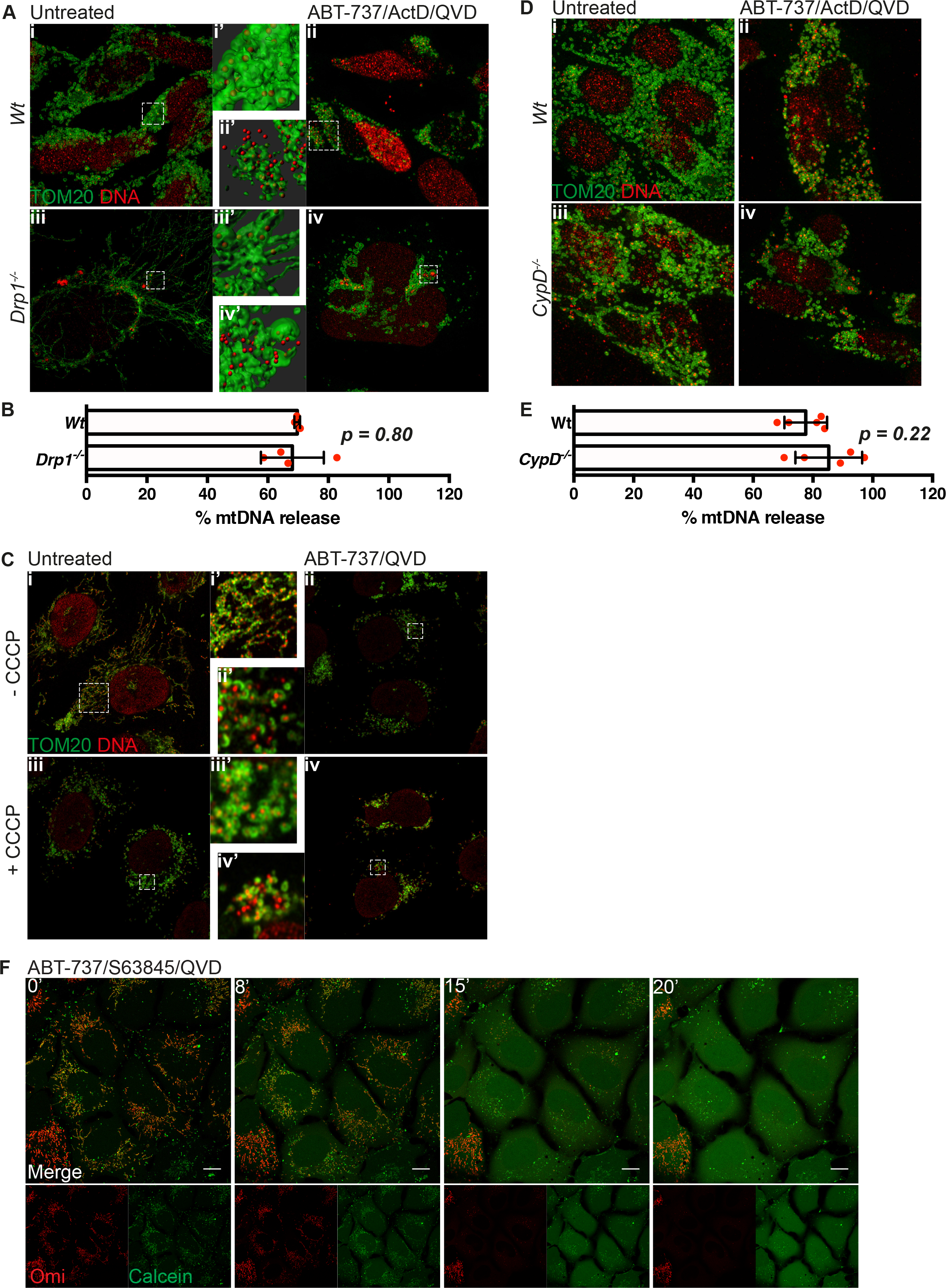
(**A**) Maximum intensity projection of z-stack Airyscan images of Drp1^*fl/fl*^ MEFs with induced Drp1 deletion by AdCre virus. Cells were treated with 10μM ABT-737, 1μM ActD and 20μM qVD-OPh for 3h and immunostained for TOM20 (MOM) and DNA. Zooms show Imaris 3D reconstructions of surface (MOM, TOM20) and spots (DNA) to show the extent of mtDNA release. Scale bar: 10μm. (**B**) Quantification of mtDNA released in (A). Z-stack Airyscan images were 3D rendered into surfaces (TOM20, MOM) and spots (DNA) and mtDNA nucleoid localization was assayed by eye. (**C**) U2OS MCL1^CRISPR^ cells were pre-treated with 10μM CCCP and 20μM qVD-OPh for 30 min to induce mitochondrial fragmentation and then treatment was changed to 10μM ABT-737 and 20μM qVD-OPh. After 3h, cells were fixed and immunostained for TOM20 and DNA. Zooms show highlighted areas. Scale bar: 10μm. (**D**) Maximum intensity projections of z-stack Airyscan images of wild-type and cyclophillin D knock-out MEFs. Cells were treated with 10μM ABT-737, 1μM ActD and 20μM qVD-OPh for 3h. (**E**) Quantification of mtDNA released in (E). Z-stack Airyscan images were 3D rendered into surfaces (TOM20, MOM) and spots (DNA) and mtDNA nucleoid localization was assayed by eye. (**F**) Representative images from time-lapse live-cell imaging of U2OS cells loaded with calcein and CoCl_2_ and treated with 10μM ABT-737/S63845/qVD-OPh at t=0. Scale bar: 10μm. (**A-C, F**) are representative data from n≥2 independent experiments. Data from (**B-E**) are compiled from cells spanning at least 2 independent experiments. Error bars represent SD.

Importantly, induced mitochondrial fission had no effect on MOMP-induced mtDNA release, consistent with the lack of requirement for DRP-1. To further define MIMP, we adapted an assay measure mitochondrial release of matrix calcein relative to MOMP by live-cell imaging (Bonora, Morganti et al., 2016). We reasoned that should the inner membrane become more permeable, Co^2+^ would be free to enter the matrix and quench matrix-located calcein, enabling of detection of increased permeability in real-time. To simultaneously visualise MOMP, we expressed Omi-mCherry which resides in the mitochondrial inner membrane space (Tait et al., 2010). U2OS cells loaded with calcein-AM and Co^2+^, were treated with ABT-737/S63845 in the presence of qVD-OPh and imaged by confocal microscopy (Figure 3D, EV Figure 4E **and Video 7**). In healthy cells, mitochondrial localised calcein persisted, consistent with the mitochondrial IMM being impermeable (**EV Figure 4D**).

**Figure 4.**
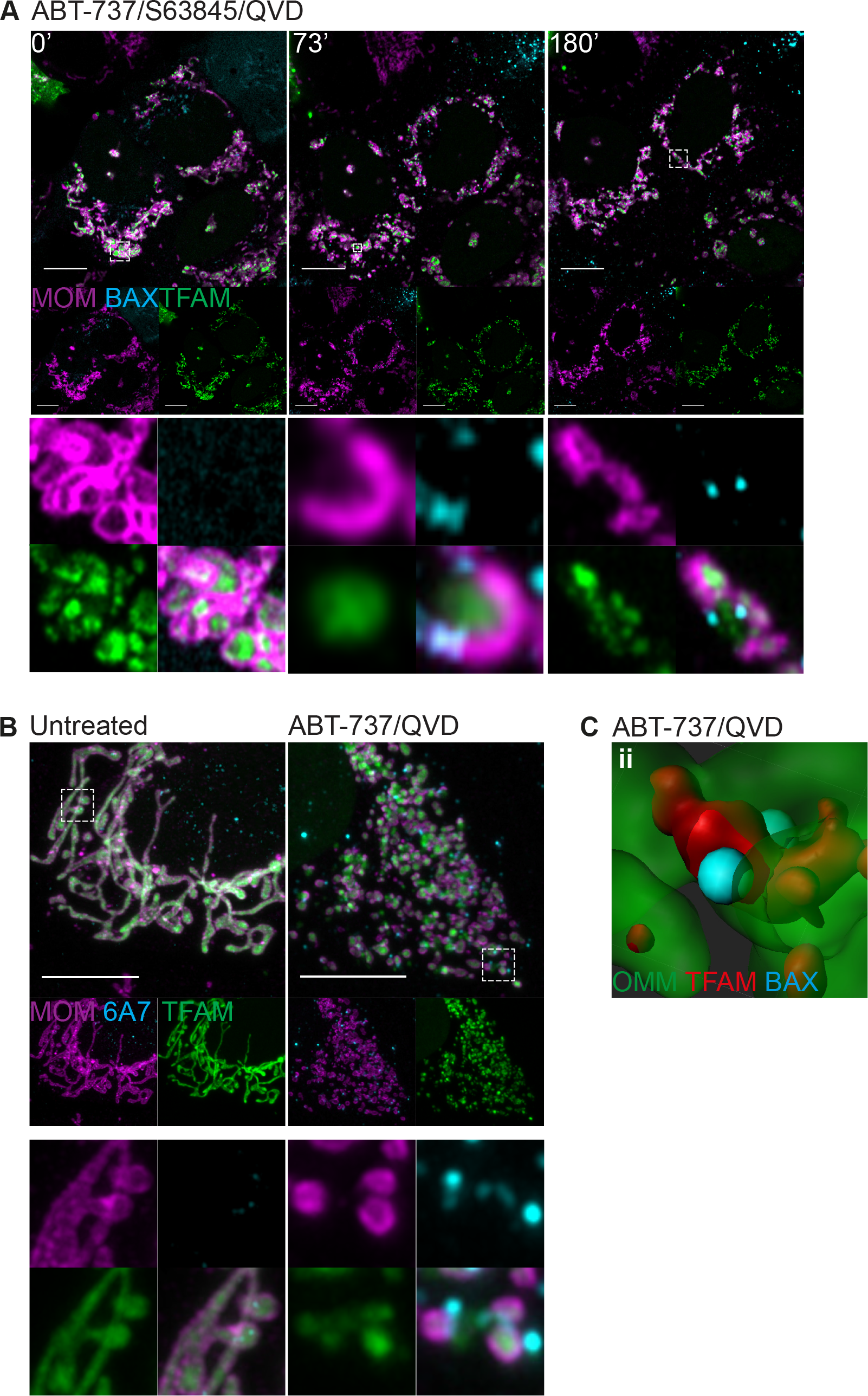
(**A**) Live-cell Airyscan imaging of U2OS BAX/BAK^CRISPR^ cells stably expressing JF_646_-MOM (magenta) and mCherry-BAX (cyan) and transiently transfected with TFAM-mClover (green). Cells were treated with 10μM ABT-737, 2μM S63845 and 20μM qVD-OPh at t=0. Scale bar: 10μm. (**B**) Maximum intensity projections of fixed cell Airyscan z-stack images of U2OS MCL1^CRISPR^ cells stably expressing SNAP-MOM (magenta) and transiently transfected with TFAM-mClover (green). Cells were treated with 2μM ABT-737 and 20μM qVD-OPh for 3h. Prior to treatment, cells were stained with JF_646_ to visualise MOM (magenta) and post fixation immunostained for active BAX (6A7, cyan). Scale bar: 10μm. (**C**) Imaris 3D reconstruction of Airyscan z-stack data of cells from (B) to show active BAX localization (cyan) in relation to TFAM-mClover (red) and MOM (green). (**A-C**) are representative of n≥2 independent experiments.

However, following MOMP (defined by mitochondrial release of Omi-mCherry) the matrix calcein signal was lost rapidly within minutes following MOMP (Figure 3D **and Video 7**). Loss of calcein signal was not blocked by co-treatment with the cyclophilin D inhibitor cyclosporin A (CsA) (**EV Figure 4E** **and Video 8**). This demonstrates, at early time points following MOMP, the mitochondrial inner membrane increases permeability to small ions. Prolonged opening of the mitochondrial permeability transition pore (MPTP) causes mitochondrial swelling leading to mitochondria rupture (Izzo, Bravo-San Pedro et al., 2016). We therefore investigated whether MPTP contributed to mtDNA release. Wild-type and MEFs deleted for cyclophilin D (EV Figure 4C) (a critical component of the MPTP)(Baines, Kaiser et al., 2005, Nakagawa, Shimizu et al., 2005)), were treated with ABT-737/ActD/qVD-OPh to engage MOMP and then immunostained for DNA and TOM20, to visualise mtDNA and the mitochondrial outer membrane respectively. Importantly, a similar extent of mtDNA release was observed between WT and CypD^−/−^ cells, ruling out a critical role for MPTP in mtDNA release (**Figures 3E** and **3F**). Collectively, these data show that MIMP, enabling the release of mtDNA, is independent of both mitochondria dynamics and MPTP.

### Mitochondrial inner membrane is extruded through expanding BAX pores

We next sought to understand how the inner membrane breaches the outer membrane, permitting mtDNA release. It is well established that during apoptosis BAX and BAK form oligomers on the mitochondrial outer membrane leading to MOMP (Cosentino & Garcia-Saez, 2017). We therefore investigated the relationship between BAX activation, MOMP and MIMP through live-cell imaging. To this end, mCherry-BAX was expressed in BAX/BAK-deleted U2OS cells together with JF_646_-MOM and TFAM-mClover. Cells were treated with ABT-737/S63845 in the presence of qVD-OPh to induce MOMP then analysed by live-cell imaging (**Figure 4A, EV Figure 5A, Video 9**). Consistent with our earlier data (**Figure 1E**), following treatment, pores in the MOM occurred that gradually increased in size (**Figures 4A, EV Figure 5A, Video 9**). Combined analysis of the MOM, TFAM and BAX showed that TFAM is commonly extruded from large MOM openings that are decorated by BAX puncta. This suggests, that growing BAX-mediated pores in the mitochondrial outer membrane enables extrusion of the mitochondrial inner membrane. To investigate this further we treated U2OS MCL1^CRISPR^ cells (which die in a rapid and synchronous manner in response to ABT-737) expressing JF_646_-MOM and TFAM-mClover with ABT-737 in the presence of qVD-OPh and immunostained with antibody recognising active BAX (6A7)(Hsu & Youle, 1997). In line with previous findings, we found that during MOMP, the MOM becomes decorated in activated BAX punctate structures (**Figure 4B, EV Figure 5B**). 3D-images were generated from Z-stacks of super-resolution images (**Figure 4C and EV Figure 5C)**. These images frequently showed translocation of TFAM-mClover through the mitochondrial outer membrane at sites demarked by active BAX staining. These data support a model, whereby following MOMP the mitochondrial inner membrane can be extruded through BAX-mediated outer membrane pores. Once in the cytoplasm the inner membrane can permeabilise, allowing mtDNA to activate cGAS/STING signaling.

## Discussion

Following apoptotic mitochondrial permeabilisation, mtDNA activates cGAS/STING-dependent interferon synthesis under caspase-inhibited conditions (Giampazolias et al., 2017, Rongvaux et al., 2014, White et al., 2014). Here we investigated how matrix localised mtDNA could activate cytosolic cGAS-STING signalling, given that their separate cellular localisations. We find that under caspase-inhibited conditions, BAX/BAK-mediated MOMP allows extrusion of the mitochondrial inner membrane into the cytoplasm. Unexpectedly, the extruded inner membrane eventually permeabilises leading to cytosolic release of mtDNA where it can activate cGAS/STING signalling. By doing so, mitochondrial inner membrane permeabilisation or MIMP, supports the immunogenic effects of caspase-independent cell death.

Surprisingly, our data shows that mitochondrial inner membrane can permeabilise subsequent to BAX/BAK-mediated MOMP. We found that over time, following MOMP (defined by release of Omi-mCherry), visible and widening pores appeared in the MOM through which the inner membrane extruded (**Figure 1E, 2B, EV Figure 3B**). Interestingly, active BAX was found to localise at the edge of the widening MOM pores through which the inner membrane extrudes (**Figure 4A, 4B**). This is reminiscent of recent studies showing that during apoptosis, BAX-rings border visible pores in the mitochondrial outer membrane (Grosse, Wurm et al., 2016, Salvador-Gallego, Mund et al., 2016). Over time we observed these MOM pores gradually widen enabling inner membrane extrusion. Coupling our data to recent findings demonstrating variation in BAX-ring size (Salvador-Gallego et al., 2016) as well as wide variation in BAX-mediated pore size (Czabotar, Westphal et al., 2013, Dewson, Ma et al., 2012) strongly suggests that BAX-mediated pores are dynamic and grow over time.

How the inner membrane eventually permeabilises remains unclear. Our data argue against any role for mitochondrial permeability transition or mitochondrial dynamics in the process. Inner membrane permeabilisation occurs considerably later (1 hour) than MOMP, with less penetrance and in apparent stochastic manner. As such, whether MIMP occurs in a regulated manner is unknown. Interestingly, we detected increased permeability of the mitochondrial inner membrane to small ions, shortly after the onset of MOMP. This increased permeability may explain the transient drop in inner membrane potential immediately following MOMP previously reported by others (Waterhouse et al., 2001). What drives this increased permeability is unknown but we reason it must be transient, since many studies have shown mitochondrial inner membrane potential can be maintained post-MOMP under caspase-inhibited conditions (Bossy-Wetzel, Newmeyer et al., 1998, Huber, Dussmann et al., 2011, von Ahsen et al., 2000, Waterhouse et al., 2001). While inner membrane extrusion and MIMP can be temporally separated from mtDNA release, it is possible that the increased permeability of the inner membrane we observe following MOMP may precipitate these later events.

Following MOMP, our ability to detect inner membrane extrusion and MIMP during cell death was dependent upon inhibition of caspase function. Purely from a technical perspective, our inability to visualise MIMP during apoptosis relates to the kinetics of apoptotic (caspase-dependent) cell death being much more rapid than MIMP, such that rounding and detachment of apoptotic cells made them impossible to image over a longer-period (**Video 1**). Even if MIMP does occur during apoptosis, it’s functional effects may be limited due rapid loss of cell viability coupled to the strong inhibitory effects of caspase activity on protein translation (Clemens, Bushell et al., 2000). Indeed, we and others have shown that MOMP has pro-inflammatory effects only under caspase-inhibited conditions (Giampazolias et al., 2017, Rongvaux et al., 2014, White et al., 2014). Nevertheless, physiological engagement of MIMP, leading to mtDNA release and interferon synthesis might be expected in cell types deficient or defective in caspase activity such as cardiomyocytes and sympathetic neurons following MOMP (Potts, Vaughn et al., 2005, Wright, Vaughn et al., 2007). Another possibility relates to our recent finding sub-lethal stress can engage MOMP in a limited cohort of mitochondria without cell death (Ichim, Lopez et al., 2015). Whether MIMP and mtDNA release occurs in these permeabilised mitochondria leading to cGAS/STING activation remains unclear but is being investigated further.

Intense interest currently surrounds the pharmaceutical activation of cGAS/STING signaling to improve cancer immunotherapy (Mullard, 2018). Our own work has shown that engaging MOMP under caspase-inhibited conditions - caspase-independent cell death (CICD) - can potently activate anti-tumour immunity (Giampazolias et al., 2017). The inflammatory effects of CICD are dependent on recognition of mtDNA by the cytosolic cGAS-STING sensing pathway (Giampazolias et al., 2017, Rongvaux et al., 2014, White et al., 2014). Underlying this effect, here we have show that following MOMP, the inner membrane is extruded via BAX-mediated pores prior to permeabilisation and mtDNA release. mtDNA dependent activation of cGAS/STING has recently been implicated in variety of pathophysiological processes including infectious and inflammatory diseases (West & Shadel, 2017). As such, further understanding the mechanism(s) of mtDNA mitochondrial release in different contexts may provide new avenues for therapeutic exploitation.

## Acknowledgments

Funding for this work was from the BBSRC (grant BB/K008374/1). We thank Luke Lavis (HHMI/Janelia Research Campus), Mikhail Alexyev (University of South Alabama), David Sabatini (MIT) for reagents. We also thank Margaret O’Prey, David Strachan and Tom Gilbey (Beatson Institute) for excellent technical assistance and members of the Tait lab for critical reading of the manuscript.

## Materials and Methods

### Cell lines, plasmids, reagents and antibodies

U2OS, MEF and 293FT were cultured in DMEM high-glucose medium supplemented with 10% FCS, 2mM glutamine, 1mM sodium pyruvate, 50μM β-mercaptoethanol, penicillin (10,000 units/mL) and streptomycin (10,000 units/mL). Reagents used were as followed: ABT-737, Q-VD-OPh (APEX Bio), S63845 (Active Biochem) carbonyl cyanide 3-chlorophenylhydrazone (CCCP), cyclosporin A (Sigma), actinomycin D (Merck). Cell lines were routinely tested for the presence of mycoplasma. No cell lines were authenticated. To delete Drp1 from Drp1^*fl/fl*^ MEFs, 2×10^6^ cells were seeded and infected with 200 MOI high titer Ad5CMVCre (Viral Vector Core, University of Iowa) for 8h, after which time the media was removed and replaced. Cells were seeded the following day for experiments. For CRISPR/Cas9-based genome editing, the following sequences were cloned into LentiCRISPRv2-puro (Addgene #52961) or LentiCRISPRv2-blasti (Lopez, Bessou et al., 2016): Human BAX: 5’- AGTAGAAAAGGGCGACAACC-3’. Human BAK: 5’- GCCATGCTGGTAGACGTGTA-3’. Human MCL1: 5’-GGGTAGTGACCCGTCCGTAC-3’. For lentiviral and retroviral transduction, the following plasmids were used: pLJM2 SNAP-Omp25 was obtained from Addgene (#69599). pcDNA3 TFAM-mClover3 was cloned using Gibson assembly (New England Biolabs) using TFAM-GFP (kind gift from Mikhail Alexeyev, University of South Alabama, (Pastukh, Shokolenko et al., 2007) and pKanCMV-mclover3-18aa-actin (Addgene #74259) as templates. mCherry-BAX was cloned into the pMx vector with blasticidin resistance using Gibson assembly. Primary antibodies for Western blotting were as follows: rabbit anti-BAX, rabbit anti-BAK, rabbit anti-DRP1 (Cell Signaling), mouse MitoProfile Membrane Integrity Cocktail (for cyclophillin D) (Abcam) and β-actin (Sigma). Primary antibodies for immunofluorescence were as follows: rabbit anti-TOM20 (Santa Cruz), mouse anti-DNA (Progen), rabbit anti-AIF (Cell Signaling), mouse anti-cytochrome *c* (BD Transduction Laboratories), mouse anti-BAX (6A7, Santa Cruz) and Alexa Fluor 647 anti-cytochrome *c* (6H2.B4, BioLegend).

### Stable cell line generation

For retroviral transduction, 293FT cells were transfected with 5μg of selected plasmid in combination with 1.2μg gag/pol (Addgene #14887) and 2.4μg UVSVG (Addgene #12260) using Lipofectamine 2000 (Life Technologies). Viral supernatant was collected, filtered and used to infect cells 24h and 48h post-transfection in the presence of 1μg/mL Polybrene (Sigma-Aldrich). Cells were selected by addition of 1μg/mL puromycin (Sigma) or 10μg/mL blasticidin (InvivoGen) as appropriate. For cell lines stably overexpressing fluorescent proteins, cells were FACS sorted to isolate a high-expressing population.

For lentiviral transduction, 293FT cells were transfected with 5 μg of selected plasmid in combination with 1.86μg psPAX2 (Addgene #8454) and 1μg UVSVG (Addgene #12260) using Lipofectamine 2000. Viral supernatant was collected, filtered and used to infect cells 24h and 48h post-transfection in the presence of 1μg/mL Polybrene. Cells were selected by addition of 1μg/mL puromycin. For cell lines stably overexpressing fluorescent proteins, cells were FACS sorted to isolate a high-expressing population.

### Microscopy

Cells were fixed in 4% PFA/PBS for 10 min, followed by permeabilisation in 0.2% Triton-X-100/PBS for 15 mins, followed by blocking in 2% BSA/PBS for 1h. Cells were incubated overnight in a humidified chamber at 4°C. Secondary antibodies used for detection were: Alexa Fluor 488 goat anti-mouse (A11029), Alexa Fluor 488 goat anti-rabbit (A11034), Alexa Fluor 568 goat anti-mouse (A11004), Alexa Fluor 568 goat anti-rabbit (A11011), Alexa Fluor 647 goat anti-mouse (A21236), Alexa Fluor 647 goat anti-rabbit (A21245); all purchased from Life Technologies. Slides were mounted in Vectashield antifade aqueous mounting medium (Vector Laboratories), with or without DAPI. For triple colour antibody-based immunofluorescence, cells were co-incubated with two antibodies (typically TOM20 and DNA), thoroughly washed and incubated with Alexa Fluor 488/568 secondary antibodies. Following extensive washing, cells were blocked with 5% mouse serum (Life Technologies) for 1h at room temperature and further incubated with cytochrome *c* directly conjugated to Alexa Fluor 647 (clone 6H2.B4, Biolegend).

#### Airyscan imaging

Super-resolution Airyscan images were acquired on a Zeiss LSM 880 with Airyscan microscope (Carl Zeiss). Samples were prepared on high precision cover-glass (Zeiss, Germany). Data were collected using a 63× 1.4 NA objective for the majority of experiments, except for calcein-AM experiments where the 40× (1.3 NA) objective was used. 488, 561 and 640nm laser lines were used and refractive index matched immersion oil (Zeiss) was used for all experiments. Where z-stacks were collected, the software-recommended optimal slice sizes were used. For both fixed-cell and live-cell experiments, colours were collected sequentially in order to minimise bleedthrough. Live cell experiments were performed in an environmental chamber to allow maintenance of 37°C temperature and 5% CO_2_. Airyscan processing was performed using the Airyscan processing function in the ZEN software. To maintain clarity and uniformity throughout the paper, some images have been pseudocoloured.

#### 3D-SIM imaging

3D structured illumination microscopy (3D-SIM) images were acquired on a Nikon N-SIM microscope (Nikon Instruments, UK). Samples were prepared on high precision cover-glass (Zeiss, Germany). Data were collected using a Nikon Plan Apo TIRF 100x 1.49 NA objective and an Andor DU-897X-5254 camera using 488, 561 and 640nm laser lines. Refractive index matched immersion oil (Nikon Instruments) were used for all experiments. Z-stacks were collected with a step size of 120nm as required by the manufacturers software. For each focal plane, 15 images were acquired (5 phases, 3 angles) and captured using the NiS-Elements software. SIM image processing, reconstruction and analysis were carried out using the N-SIM module of the NiS-Elements Advanced Research software. Images were checked for artefacts using the SIMcheck software (http://www.micron.ox.ac.uk/software/SIMCheck.php). Images were reconstructed using NiS Elements software v4.6 (Nikon Instruments, Japan) from a z-stack comprising of no less than 1um of optical sections. In all SIM image reconstructions the Wiener and Apodization filter parameters were kept constant

#### 3D renderings and mtDNA release quantification

Acquired z-stacks were imported into Imaris (Bitplane, Oxford Instruments, Switzerland). To segment inner mitochondrial membrane (IMM) and mtDNA nucleoids, a surface was creating using the IMM channel. Masks were applied to differentiate between mtDNA nucleoids inside and outside the surface (i.e. in or out of the IMM). From these masks, spots were created from the mtDNA channel.

For quantification of mtDNA outside the mitochondrial outer mitochondrial (MOM) a surface was created for the MOM channel. Spots were then generated from the mtDNA channel and the transparency of the MOM surface altered to allow visualisation of mtDNA nucleoid spots outside or inside the MOM. Cells were visually scored to quantitate the degree of mtDNA release.

#### SNAP-tag labelling

Cell lines stably expressing pLJM2 SNAP-Omp25 were incubated with 15nM JF_646_ SNAP ligand (kind gift from Luke Lavis) for 30 mins in complete medium. Throughout the paper we refer to MOM labelled in this manner as JF_646_-MOM. Cells were washed 3 times in medium and returned to the incubator for 15 mins to allow any unbound dye to diffuse out. Media was replaced with FluoroBrite DMEM (Life Technologies) supplemented with 10% FCS, 2mM glutamine and 1mM sodium pyruvate and then imaged.

### Calcein-AM assay

Mitochondria inner membrane permeabilisation was assessed using a modified version of the calcein/cobalt assay described by Bonora et al (PMID: 27172167). Briefly, U2OS cells stably expressing JF_646_-MOM and Omi-mCherry were cells seeded onto coverglass dishes to 70% confluency. Cells were incubated with 1μM of calcein stock solution and 2mM CoCl_2_ in HBSS for 15 min at 37°C in a 5% CO_2_ atmosphere. Cells were washed twice in HBSS and then 900μL of HBSS was added. Cells were left to acclimatise in the microscope environmental chamber for 30 min and imaged following addition of inducers of apoptosis, qVD-OPh ± cyclosporin A.

### Western blotting

Cells were lysed in NP-40 lysis buffer (1% NP-40, 1mM EDTA, 150mM NaCl, 50mM Tris-Cl pH7.4) supplemented with complete protease inhibitor (Roche). Protein concentration was determined using the Bradford assay (BioRad). Protein lysates were subjected to electrophoresis through 10% or 12% SDS-PAGE gels and transferred onto nitrocellulose membrane. Membranes were blocked in 5% milk/PBS-Tween for 1h at room temperature and incubated with primary antibody over night at 4°C in blocking buffer. Following washing, membranes were incubated with Li-Cor secondary antibodies (IRDye 680RD donkey anti-mouse or IRDye 800CW donkey anti-rabbit) for 1h at room temperature and protein levels detected using a Li-Cor Odyssey CLx (Li-Cor). Resulting images were minimally processed for clarity and arranged using Adobe Illustrator.

### Live-cell viability assays

Cell viability was assayed using an IncuCyte FLR imaging system (Sartorius). Briefly, cells were seeded and drugged in the presence of 30nM SYTOX Green (Life Technologies), a non-cell-permeable nuclear stain. Where the same cell line was compared, SYTOX Green-positivity was quantified directly; where different cell lines (for example, WT and KO MEFs) were compared, data were normalised to confluency. All quantification was performed using the IncuCyte software.

## Video 1

U2OS cells stably expressing JF_646_-MOM (magenta) and Omi-mCherry (red) and transiently expressing TFAM-mClover (green) were treated with 10μM ABT-737 and 2μM S63845.

## Video 2

U2OS cells stably expressing JF_646_-MOM (red) and transiently expressing TFAM-mClover (green) were treated with 10μM ABT-737 and 2μM S63845 in the presence of 20μM qVD-OPh. Scale bar: 10μm.

## Video 3

Zoom of **Video 2**. Scale bar: 10μm.

## Video 4

U2OS cells stably expressing JF_646_-MOM (magenta), Omi-mCherry (red) and transiently expressing TFAM-mClover (green) were treated with 10μM ABT-737 and 2μM S63845 in the presence of 20μM qVD-OPh.

## Video 5

3D view of Imaris reconstructions of untreated U2OS cells immunostained for MIM (AIF) and DNA. mtDNA outside (cyan) and inside the MIM (green) were quantified.

## Video 6

3D view of Imaris reconstructions of U2OS cells treated with 10μM ABT-737, 1μM ActD and 20μM qVD-OPh immunostained for MIM (AIF) and DNA. mtDNA outside (cyan) and inside the MIM (green) were quantified.

## Video 7

U2OS cells stably expressing JF_646_-MOM and Omi-mCherry loaded with calcein and CoCl_2_ were treated with 10μM ABT-737, 2μM S63845 and 20μM qVD-OPh and imaged.

## Video 8

U2OS cells stably expressing JF_646_-MOM and Omi-mCherry loaded with calcein and CoCl_2_ were treated with 10μM ABT-737, 2μM S63845 and 20μM qVD-OPh in the presence of 25μM cyclosporin A and imaged.

## Video 9

U2OS cells stably expressing JF_646_-MOM and BAX-mCherry and TFAM-mClover were treated with 10μM ABT-737, 2μM S63845 and 20μM qVD-OPh and imaged.

---

**EV Figure 1.**
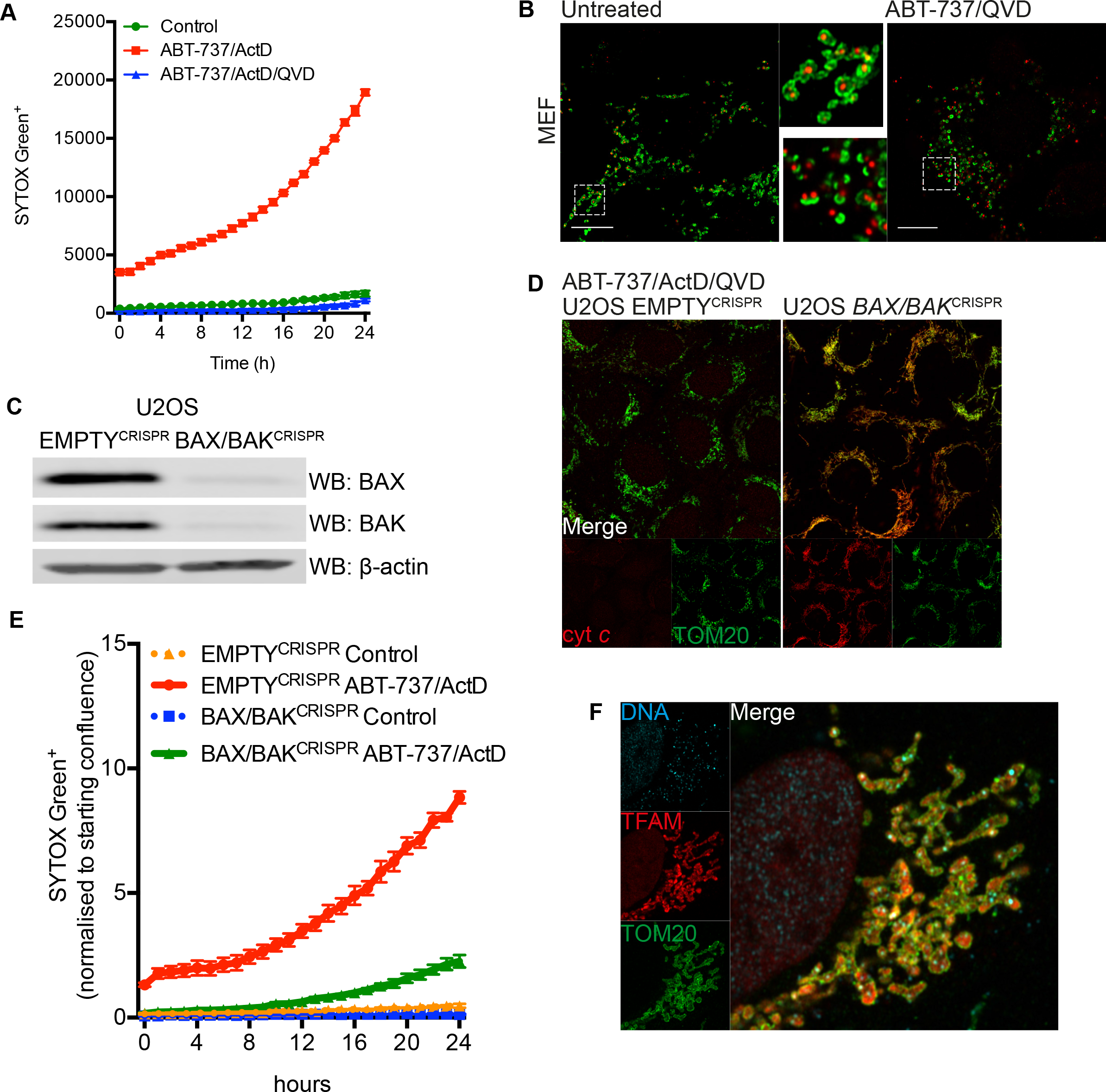
(**A**) Live-cell IncuCyte imaging of cell death measured by SYTOX Green uptake of U2OS cells treated with 10μM ABT-737, 1μM ActD ± 20μM qVD-OPh or 10μM ABT-737, 2μM S63845 ± 20μM qVD-OPh. (**B**) Wild-type MEFs treated with 10μM ABT-737 and 20μM qVD-OPh for 3h. Airyscan images of cells immunostained with anti-TOM20 and anti-DNA antibodies. Scale bar: 10μm. (**C**) Western blot analysis showing CRISPR-Cas9-mediated co-deletion of BAX and BAK in U2OS cells as assessed by immunoblotting for BAX and BAK. B-actin serves as a loading control. (**D**) Airyscan images of U2OS EMPTY^CRISPR^ and BAX/BAK^CRISPR^ cells treated with 10μM ABT-737, 1μM ActD and 20μM qVD-OPh for 3h immunostained for cytochrome c and TOM20. (**E**) Live-cell IncuCyte imaging of SYTOX Green uptake of U2OS EMPTY^CRISPR^ and BAX/BAK^CRISPR^ cells treated with 10μM ABT-737 and 1μM ActD. (**F**) Airyscan images of U2OS cells transiently transfected with TFAM-mClover (pseudocoloured in red), and immunostained with TOM20 and DNA. (**A, E**) are representative data from n≥2 independent experiments. Error bars represent SEM. (**B**) is a representative image of n≥2 independent experiments. (**C, D, F**) shows representative data from n≥2 independent experiments.

**EV Figure 2.**
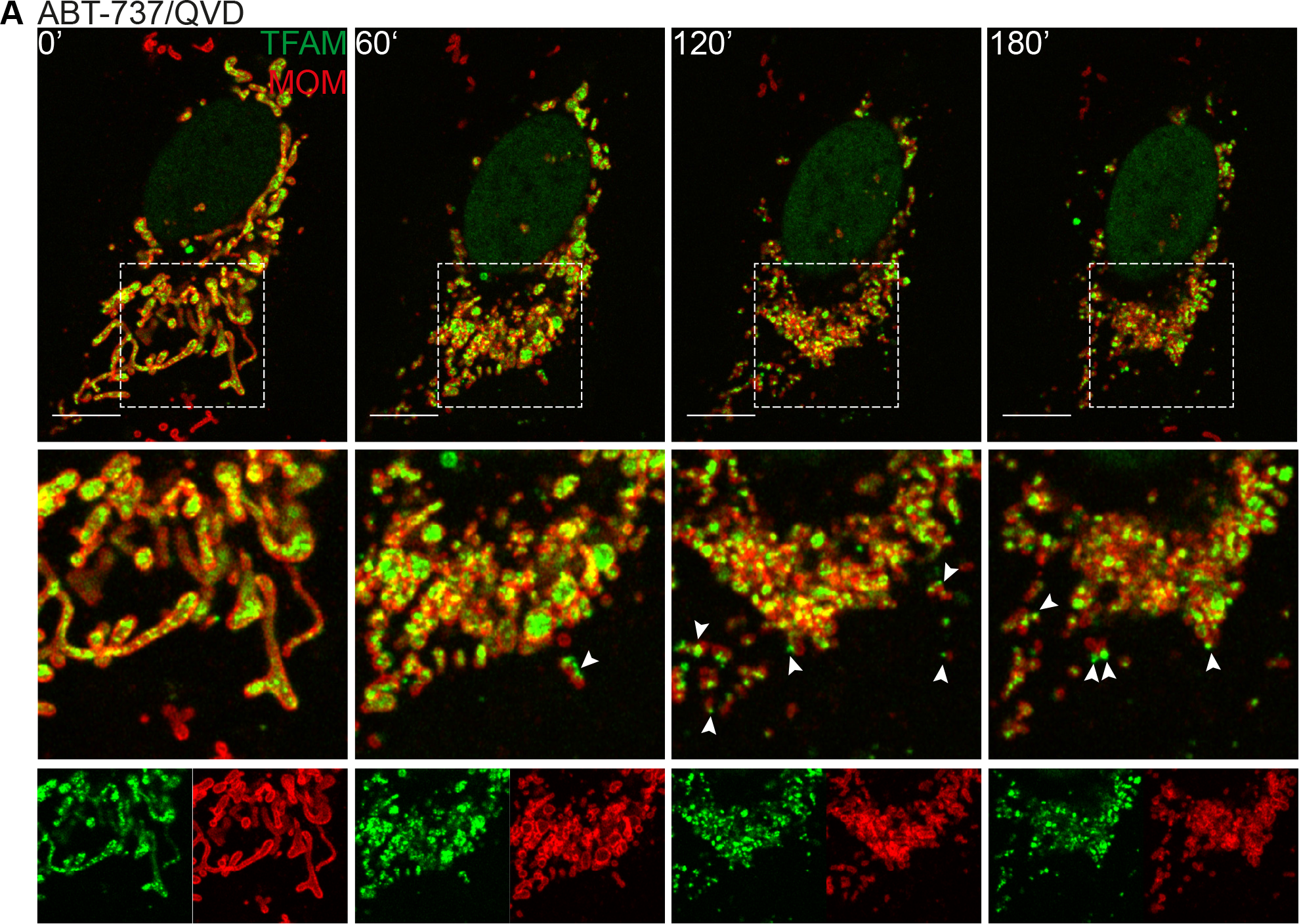
(**A**) Live-cell Airyscan imaging of U2OS *MCL1*^CRISPR^ cells stably expressing JF_646_-MOM and transiently transfected with TFAM-mClover. Cells treated with 2μM ABT-737 and 20μM qVD-OPh at t=0. Scale bar: 10μm. Data representative of n≥3 independent experiments.

**EV Figure 3.**
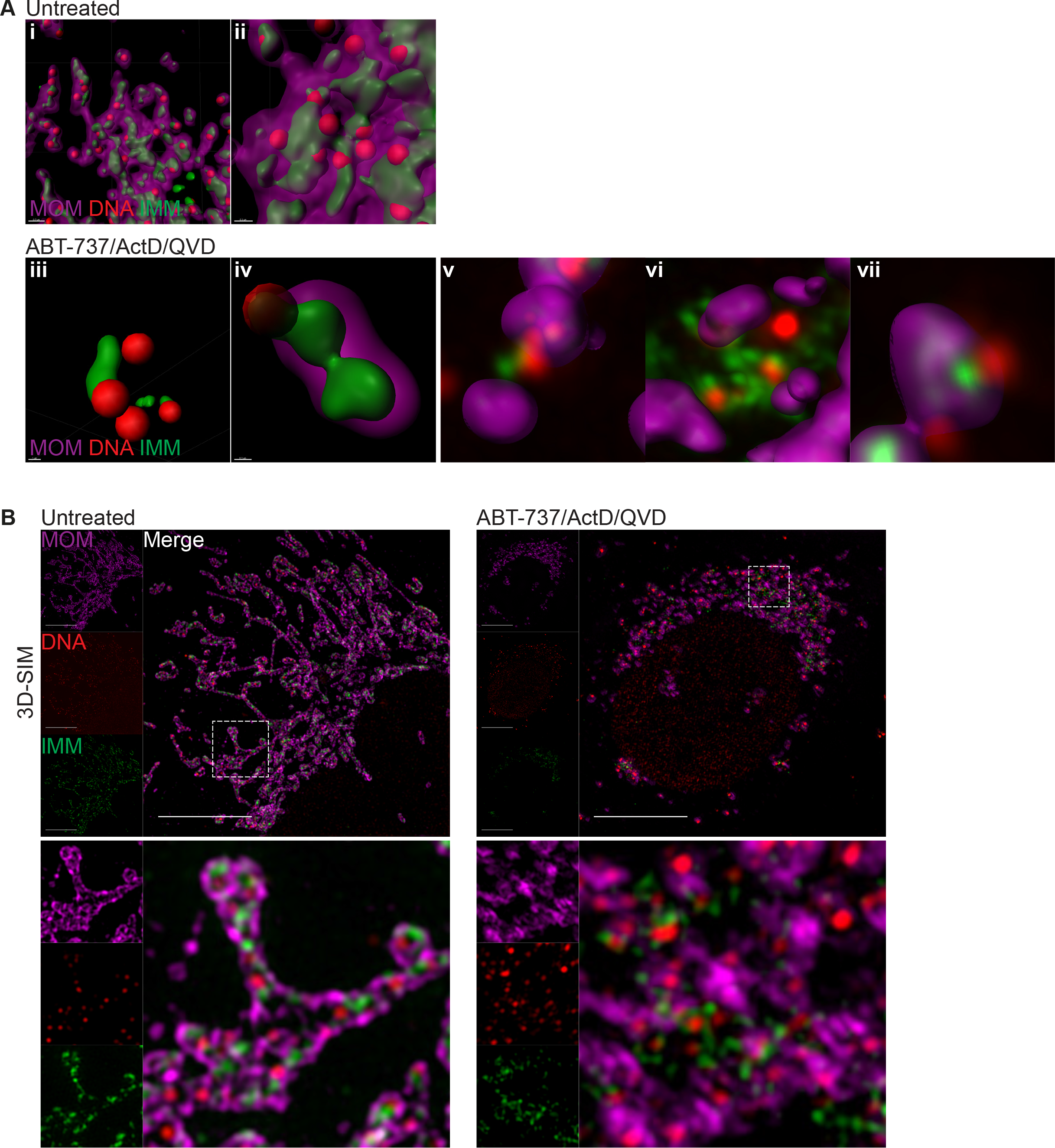
(**A**) Further examples of Imaris 3D reconstructions from **Figure 2B** of (**i-ii**) U2OS untreated cells and (**iii-vii**) cells treated with 10μM ABT-737, 1μM ActD and 20μM qVD-OPh. Scale bars as follows: (i) 0.7μm, (ii) 0.3μm, (iii) 1μm, (iv) 0.3μm. (**B**) 3D-SIM images of fixed U2OS cells treated with 10μM ABT-737, 1μM ActD and 20μM qVD-OPh for 3h. Prior to treatment cells were labelled with JF_646_ (to label SNAP-MOM)and post-fixation were immunostained for IMM (AIF) and DNA. Scale bar: 10μm. (**A-B**) are representative data from n≥3 independent experiments.

**EV Figure 4.**
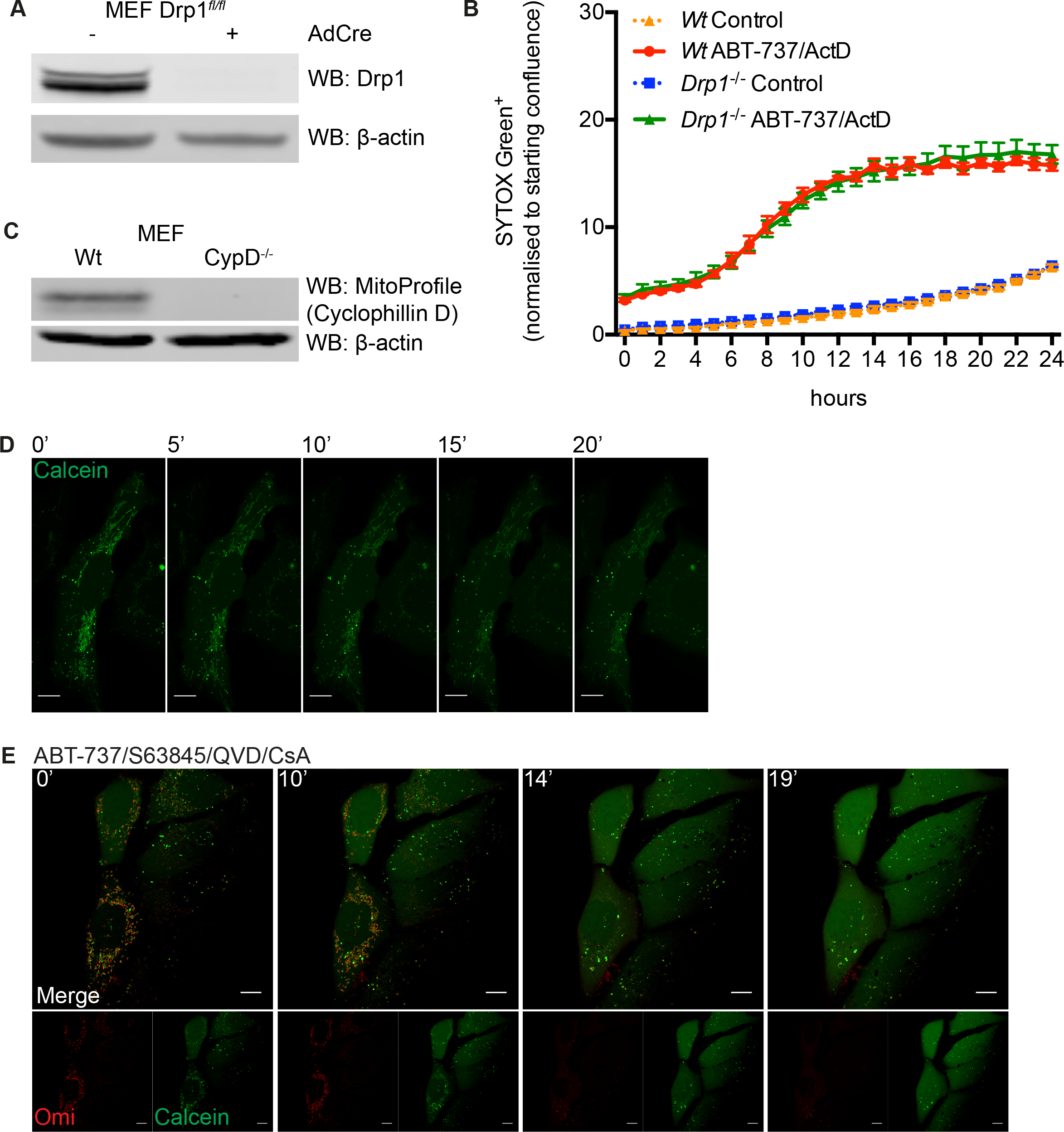
(**A**) Western blot analysis of Drp1^*fl/fl*^ MEFs ± AdCre to show deletion of Drp1. B-actin serves as a loading control. (**B**) Live-cell IncuCyte imaging of SYTOX Green uptake of Drp1^*fl/fl*^ MEFs ± AdCre cells treated with 10μM ABT-737 and 1μM ActD. (**C**) Western blot analysis of wild-type and cyclophillin D MEFs to show deletion of cyclophillin D. MitoProfile membrane integrity antibody cocktail, which contains anti-cyclophillin D was used to visualise cyclophillin D. B-actin serves as a loading control. (**D**) Representative images from time-lapse live-cell imaging of U2OS cells loaded with calcein and CoCl_2_ and treated with 10μM ABT-737/S63845/qVD-OPh in presence of 25μM CsA at t=0. Scale bar: 10μm. (**E**) U2OS cells loaded with calcein-AM and CoCl2 and imaged every 30s for 20 min to show absence of photobleaching. Scale bar: 10μm. (**A-E**) are representative of n≥2 independent experiments. Error bars in (**B**) represent SEM.

**EV Figure 5.**
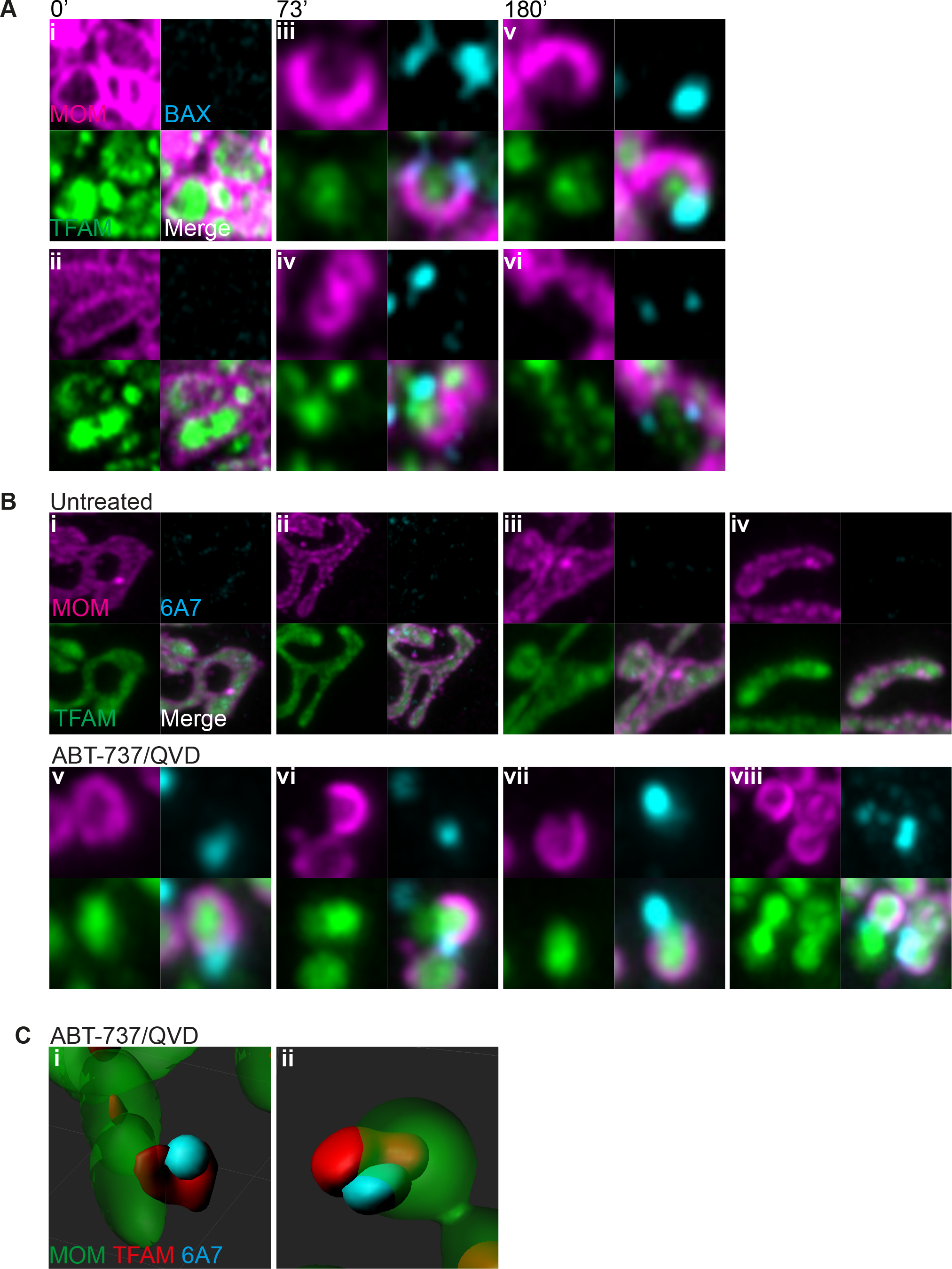
(**A**) Further examples of active BAX localization (cyan) in relation to TFAM-mClover (green) and MOM (magenta). (**B**) Further examples of Imaris 3D reconstructions of Airyscan data from Figure 4B showing active BAX (cyan), TFAM-mClover (red) and MOM (green). (**A-C**) are representative of n≥2 independent experiments.

